# A Novel Patient-Derived Mutation in Glucocorticoid Receptor Reveals an Essential Role for DNA binding in Development and Endocrine Regulation

**DOI:** 10.64898/2025.12.12.693739

**Authors:** Theophilus T. Tettey, Qizong Lao, Sonal Vaid, Sohyoung Kim, Lyuba Varticovski, Vipula Kolli, Deborah P. Merke, Gordon L. Hager

## Abstract

Glucocorticoid receptors (GRs) are vital transcription factors regulating stress responses, metabolism, and development. We report here a novel heterozygous *NR3C1* c.1310C>T (p.T437I) variant within the GR DNA-binding domain found in a family exhibiting clinical and hormonal manifestations of partial glucocorticoid resistance. Patient-derived fibroblasts demonstrated an overall suppression in dexamethasone-induced transcription. To comprehensively assess the significance of the variant, we engineered a knock-in mouse model (*Nr3c1^+/T444I^* mice) using CRISPR-Cas9, providing an *in vivo* model of the patient-derived GR mutation at the orthologous residue. Heterozygous mice exhibited partial glucocorticoid resistance, dysregulated HPA axis activity and impaired dexamethasone suppression, closely recapitulating the patient phenotype. Homozygous *Nr3c1^T444I/T444I^* embryos were recovered at embryonic day 12.5 (E12.5) but did not survive to term, indicating mid-gestational lethality. Transcriptomic profiling of primary mouse embryonic fibroblasts revealed dosage-dependent effects with heterozygotes showing an intermediate response to dexamethasone compared to wild-type, while homozygotes showed a markedly blunted response. Our results challenge prior assumptions by demonstrating that GR DNA-binding is essential for embryogenesis, while offering a new preclinical platform to investigate glucocorticoid resistance pathophysiology and therapeutic interventions.

## Introduction

The glucocorticoid receptor (GR), encoded by the human *NR3C1* gene, is a ligand-activated transcription factor that regulates a diverse array of physiological processes, including immune responses, metabolism, and stress adaptation (1–4). Upon binding glucocorticoids, such as cortisol, GR translocates to the nucleus, where its DNA-binding domain (DBD) interacts with glucocorticoid response elements (GREs) to modulate gene expression (2). Dysregulation of GR function is implicated in numerous disorders, including severe depressive disorder, where hypothalamic-pituitary-adrenal (HPA) axis hyperactivity and elevated cortisol levels suggest impaired GR-mediated feedback (5–7), and primary generalized glucocorticoid resistance, a rare condition characterized by partial or complete insensitivity to glucocorticoids, leading to compensatory HPA axis activation (8–10). Recent advancements in sequencing technology have made it increasingly possible to probe the human genome, leading to the identification of numerous *NR3C1* mutations (10–12). While many of these mutations are associated with diverse clinical phenotypes, the genotype-phenotype relationship remains elusive, with mutations across GR resulting in variable manifestations, from partial resistance to severe endocrine dysfunction (8, 10, 13–24). Animal models, particularly mice, have been instrumental in elucidating the functional consequences of GR mutations, providing valuable insights into stress responses, metabolism, and immune regulation (25–31). However, most existing mouse models harbor engineered GR mutations that do not correspond to variants identified in humans, limiting their translational relevance.

The GR DNA-binding domain (DBD), essential for GRE interaction and transcriptional regulation, is particularly critical, yet patient-derived DBD mutations and their *in vivo* significance have been underexplored (17, 32, 33). Here, we report a novel heterozygous *NR3C1* c.1310C>T variant resulting in a threonine to isoleucine at amino acid position 437 within the DBD of GR, identified in a family exhibiting partial glucocorticoid resistance, characterized by elevated cortisol, incomplete dexamethasone suppression, and metabolic abnormalities. To investigate the functional consequences of this mutation, we used CRISPR-Cas9 to generate a knock-in mouse model harboring the orthologous T444I mutation in the murine *Nr3c1* gene. Heterozygous mice recapitulated the patient’s phenotype, exhibiting partial glucocorticoid resistance and dysregulated HPA axis activity, while homozygous mutants were embryonically lethal at mid-to-late gestation, underscoring the critical role of this DBD of GR in development. This model bridges the gap between genetic findings and functional outcomes, providing a robust platform to investigate the molecular mechanisms of GR dysfunction and explore therapeutic strategies for glucocorticoid resistance syndromes.

## Results

### Case: A family with partial glucocorticoid resistance associated with a heterozygous *NR3C1* c.1310C>T (p.T437I) variant

The proband, a 14-year-old male (Fig. 1A, II.3; Fig. 1B), presented with hypertension and a history of early pubic hair with advanced bone age, consistent with mineralocorticoid and adrenal androgen excess, respectively. Additional significant medical history included chronic fatigue, developmental delay, anxiety, attention deficit hyperactivity disorder (ADHD), and asthma resistant to steroid treatment. Biochemical assessment revealed elevated adrenal androgens (dehydroepiandrosterone 813 ng/dL, normal range <500; androstenedione 209 ng/dL, normal range 50–100), and elevated 24-hour urine free cortisol (366.2 µg/24 h; normal range 4.0–56.0), elevated morning and evening adrenocorticotropic hormone (ACTH) levels without loss of diurnal cortisol variation (Fig. 1C,D), and failed cortisol suppression with 1 and 2 mg dexamethasone administration; cortisol suppressed with 3 mg dexamethasone administration, indicative of partial glucocorticoid resistance (Fig. 1E). Adrenal computed tomography (CT) revealed a 0.6 cm left adrenal nodule. His mother (Fig. 1A, I.1) had a normal hormonal evaluation and appropriate suppression of cortisol following 1 mg of dexamethasone (Fig. 1E); she exhibited no signs or symptoms of glucocorticoid resistance. His father (Fig. 1A, I.2; Fig. 1B) had normal pubertal development but had hypertension with biochemical evidence of elevated cortisol (24-hour urinary free cortisol 93.1 µg/24 hours; normal range 3.5–45), elevated morning and evening ACTH levels without loss of diurnal cortisol variation (Fig. 1C,D), and failed cortisol suppression with 1 mg dexamethasone; cortisol suppressed with 2 mg dexamethasone (Fig. 1E). Adrenal CT revealed nodular thickening of the left adrenal gland. The proband’s younger brother (Fig. 1A, II.4) had a significant history of headaches, fatigue, low exercise tolerance, and elevated blood pressures in the setting of hyporeninemia and low aldosterone, delineating increased cortisol activity at the mineralocorticoid receptors (Fig. 1B). His 24-hour urinary free cortisol was elevated (64.0 µg/24 hours; normal range 2.6–37) and morning cortisol levels did not fully suppress with 1 mg or 2 mg dexamethasone but did suppress with 3mg (Fig. 1E). Adrenal CT revealed a 0.8 cm left adrenal nodule. Clinical exome sequencing (GeneDX) identified a heterozygous c.1310C>T variant in exon 3 of the *NR3C1* gene, resulting in a threonine-to-isoleucine substitution at codon 437 (p.Thr437Ile). Sanger sequencing confirmed segregation of the variant with the phenotype in the proband, his father, and younger brother, while the unaffected mother carried the wild-type allele (Fig. 1F). The variant is absent from gnomAD (34) and was recently submitted to ClinVar by GeneDX (35). The biochemical findings in the family, together with the incomplete dexamethasone suppression and familial segregation of the novel *NR3C1* DNA-binding domain variant, support a diagnosis of partial glucocorticoid resistance associated with impaired GR-mediated feedback inhibition. Decreased sensitivity to glucocorticoids disrupts hypothalamic–pituitary negative feedback, causing compensatory ACTH hypersecretion. This, in turn, drives excess adrenal steroid production, leading to mineralocorticoid and androgenic effects as observed in this family.

**Figure 1.**
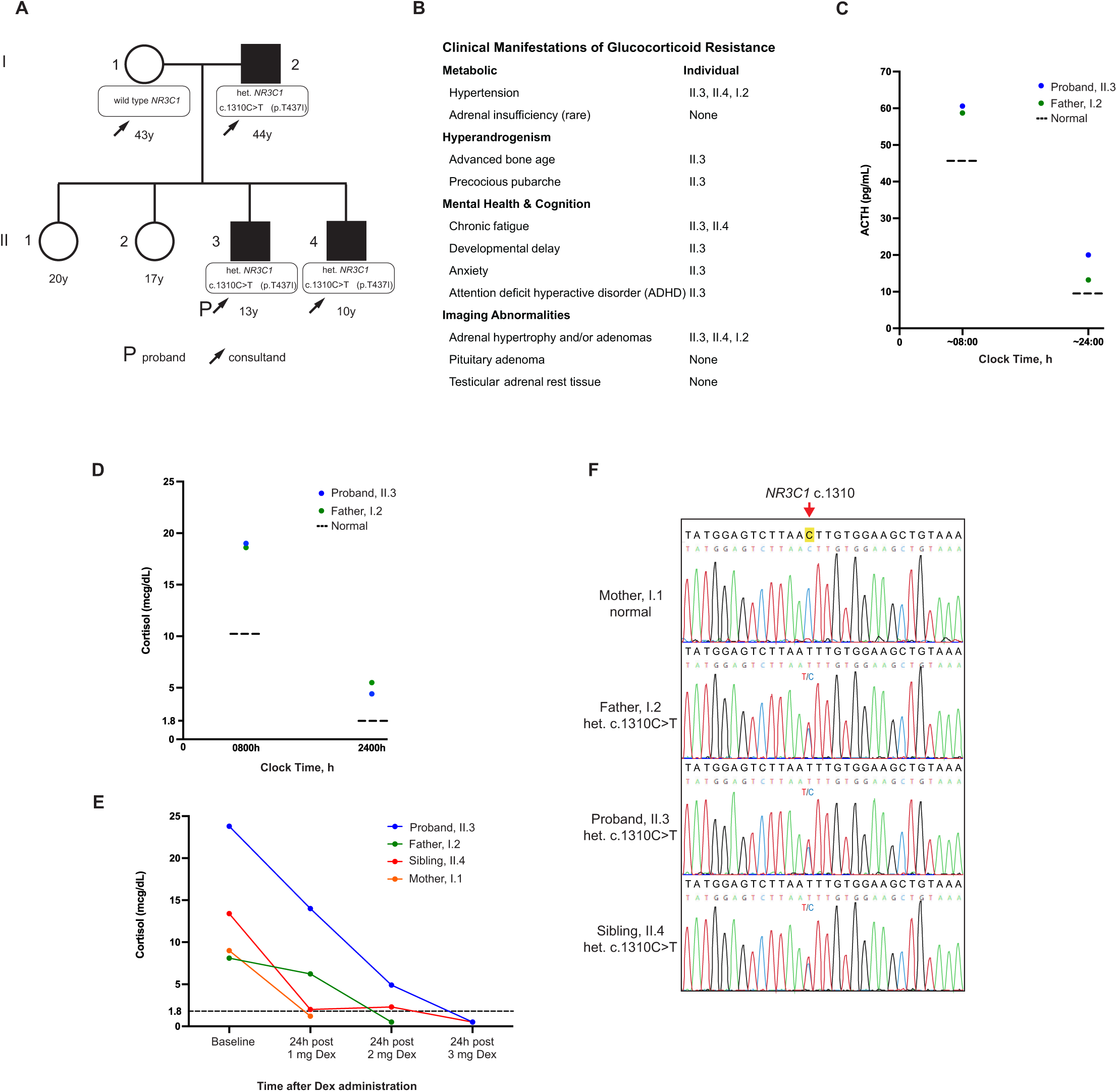
Patient characteristics associated with partial glucocorticoid resistance. (A) Pedigree of the family showing the proband (II.3), brother (II.4), and father (I.2) with partial glucocorticoid resistance and unaffected mother (I.1) and siblings (II.1, II.2) by family history. Filled symbols indicate affected individuals, open symbols indicate unaffected individuals, and genotypes are denoted where assessed. (B) Clinical features characteristic of partial glucocorticoid resistance of family members. (C) Diurnal variation of plasma ACTH measured at 0800h and 2400h, showing elevated ACTH in proband and father. Normal reference ranges are indicated by dashed lines (AM: 5–46 pg/mL; PM: <5–10 pg/mL). (D) Diurnal variation of serum cortisol measured at 0800h and 2400h. The proband and father demonstrated elevated nighttime cortisol compared to reference ranges (AM >10 μg/dL; PM <1.8 μg/dL). Dashed lines indicate normal values. (E) Dexamethasone suppression test provides cortisol levels in response to escalating doses of dexamethasone. Higher levels of dexamethasone were needed to achieve the expected cortisol suppression to <1.8 mcg/dL in the proband and father. (F) Sanger sequencing chromatograms confirming the heterozygous *NR3C1* c.1310C>T (p.Thr437Ile) variant in the proband (II.3), affected relatives, and the wild-type sequence in the unaffected mother (I.1).

### Impaired Transcriptional Response to Dexamethasone Stimulation in Patient Fibroblasts

The c.1310C>T (p.Thr437Ile) mutation is located within the DNA-binding domain (DBD) of GR, within the proximal zinc finger motif, critical for DNA binding (Fig. 2A,B). Structural analysis using PDBePISA suggests that substitution of threonine, a polar uncharged amino acid, with isoleucine, a bulkier nonpolar amino acid, at position 437, does not disrupt GR dimerization but alters the DBD’s folding dynamics, potentially impairing interaction with glucocorticoid response elements (GREs) (Fig. 2B). This structural alteration may compromise the receptor’s ability to regulate target gene expression. Multiple sequence alignment across species shows that threonine at position 437 is highly conserved in over 99% of sequences, with ConSurf analysis assigning a conservation score of 9, indicating strong evolutionary constraint (Supplementary Table 1). In silico prediction tools further support the deleterious nature of the mutation: SIFT (score: 0.00, deleterious), PolyPhen-2 (score: 1.000, probably damaging), REVEL (score: 0.932), ClinPred (score: 0.998), and MutPred2 (score: 0.838, predicting altered DNA and metal binding). A CADD PHRED-like score of 32 places the variant in the top 0.1% of deleterious substitutions. SpliceAI predicted no impact on splicing, suggesting the mutation’s effect is likely confined to the level of protein function (Supplementary Table 1). To assess the functional consequences of the mutation, we performed RNA sequencing on skin fibroblasts from the proband (II.3) and three healthy controls (1, 2, and 3) after 2 hours of treatment with 100 nM dexamethasone or vehicle (ethanol). Using a p-value < 0.05 and a fold-change ≥1.3, we identified 1,972 differentially expressed genes (DEGs), with the proband showing a significantly reduced transcriptional response compared to the controls (Fig. 2C, Extended Data Fig. 2A-C). Cluster analysis revealed three distinct patterns: Cluster C1 (671 genes) showed higher basal expression in the proband with no significant dexamethasone-induced change; Cluster C2 (682 genes) was induced by dexamethasone in controls but remained suppressed in the proband; Cluster C3 (639 genes) was repressed by dexamethasone in controls and in the proband (Fig. 2C). Gene set enrichment analysis (GSEA(36)) of HALLMARK pathways highlighted shared and distinct enrichments post-dexamethasone (Extended Data Fig. 2d). As expected, dexamethasone treatment in healthy controls led to suppression of several immune- and inflammation-related pathways, including TNFα signaling via NF-κB, inflammatory response, and allograft rejection. Similar suppression was observed in the proband, consistent with partial preservation of GR-mediated anti-inflammatory activity. By contrast, other pathways diverged: the p53 pathway was suppressed in the proband but induced in controls, while immune-related programs such as the interferon-α response, complement, and interferon-γ response were suppressed in the proband yet enriched in controls. These differences indicate impaired GR-dependent regulation of stress and immune signaling. These findings are consistent with previous reports of GR mutations causing partial glucocorticoid resistance, where impaired transcriptional activation leads to endocrine and metabolic abnormalities (8–10, 37). The T437I mutation’s impact on the DBD is consistent with a mechanistic basis for the proband’s clinical symptoms, highlighting the critical role of this conserved residue in GR function.

**Figure 2.**
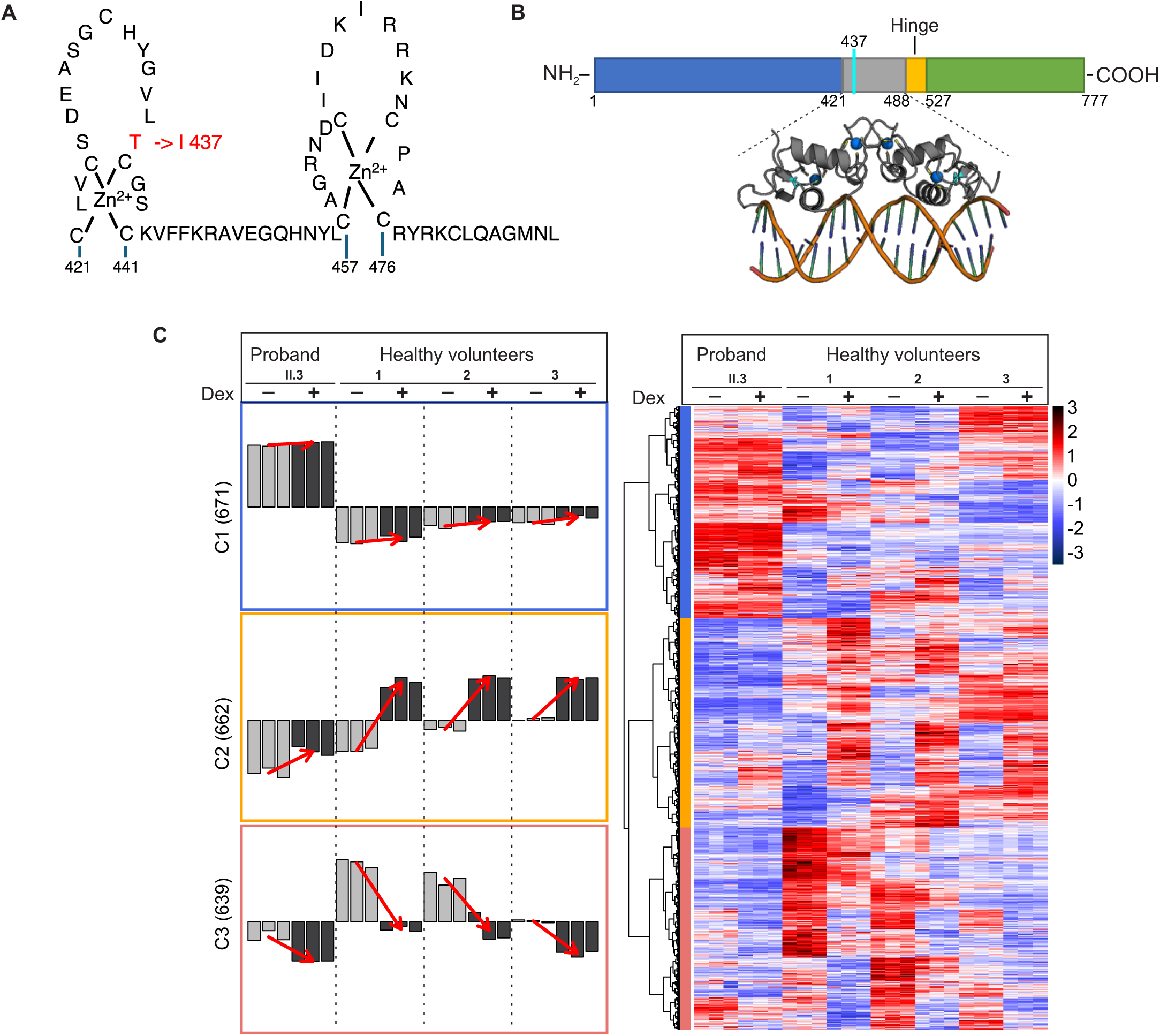
Functional consequences of the GR-T437I mutation on glucocorticoid receptor (GR)-mediated transcription in patient-derived fibroblast. (A) Schematic representation of the human glucocorticoid receptor (hGR) DNA-binding domain (DBD) (residues 421–488), highlighting the T437I mutation (red) located within the proximal zinc finger motif. (B) Domain organization of hGR, showing the N-terminal domain (NTD), DNA-binding domain (DBD), ligand-binding domain (LBD), and the hinge region. The structural model (PDB: 1R4O) illustrates the GR-DBD homodimer (gray) bound to DNA, with the T437I mutation site marked in cyan. (C) The heatmap (right panel) displays 1,972 differentially expressed genes (DEGs) in fibroblasts from the proband (II.3) compared with three healthy volunteers (1, 2, and 3) following dexamethasone (Dex, +) or vehicle (-) treatment. Hierarchical clustering of expression profiles identified three major clusters (C1, C2, C3), revealing altered Dex responsiveness in the proband. Bar plots (left panel) represent the average z-scored, variance-stabilized gene expression values for each cluster; three bars per condition indicate biological triplicates. Red arrows denote the direction of the Dex-induced response based on mean cluster expression profiles. DEGs were defined using false discovery rate (FDR) < 0.05 and |log2 fold change| > log2(1.3).

### Patient-Derived GR DBD *Nr3c1* Mouse Model Reveals Developmental Impacts

To investigate the in vivo consequences of the human GR T437I mutation, we used CRISPR–Cas9 genome editing to generate a knock-in mouse model carrying the orthologous T444I substitution in exon 3 of the *Nr3c1* gene (Fig. 3A). The T444I mutation was introduced alongside a synonymous nucleotide change that created a diagnostic *BamHI* restriction site. Founder mice were identified by restriction fragment length polymorphism (RFLP) analysis (Fig. 3C) and verified by Sanger sequencing (Fig. 3D), followed by backcrossing over multiple generations to stabilize the mutant allele and minimize off-target effects (workflow shown in Fig. 3B). Heterozygous *Nr3c1*-T444I mice (GR^+/T444I^) were viable, fertile, and exhibited no overt developmental abnormalities. Breeding with wild-type mice yielded offspring in expected Mendelian ratios (∼50% GR^+/T444I^, 50% GR^WT^; n=210, genotyped at 3 weeks; Table 1). In contrast, intercrosses between heterozygous GR^+/T444I^ mice produced no viable homozygous GR^T444I/T444I^ offspring at birth, indicating embryonic lethality (Table 1). Timed mating experiments revealed that homozygous embryos were recoverable at embryonic day 12.5 (E12.5), suggesting lethality occurs between mid- and late gestation.

**Figure 3.**
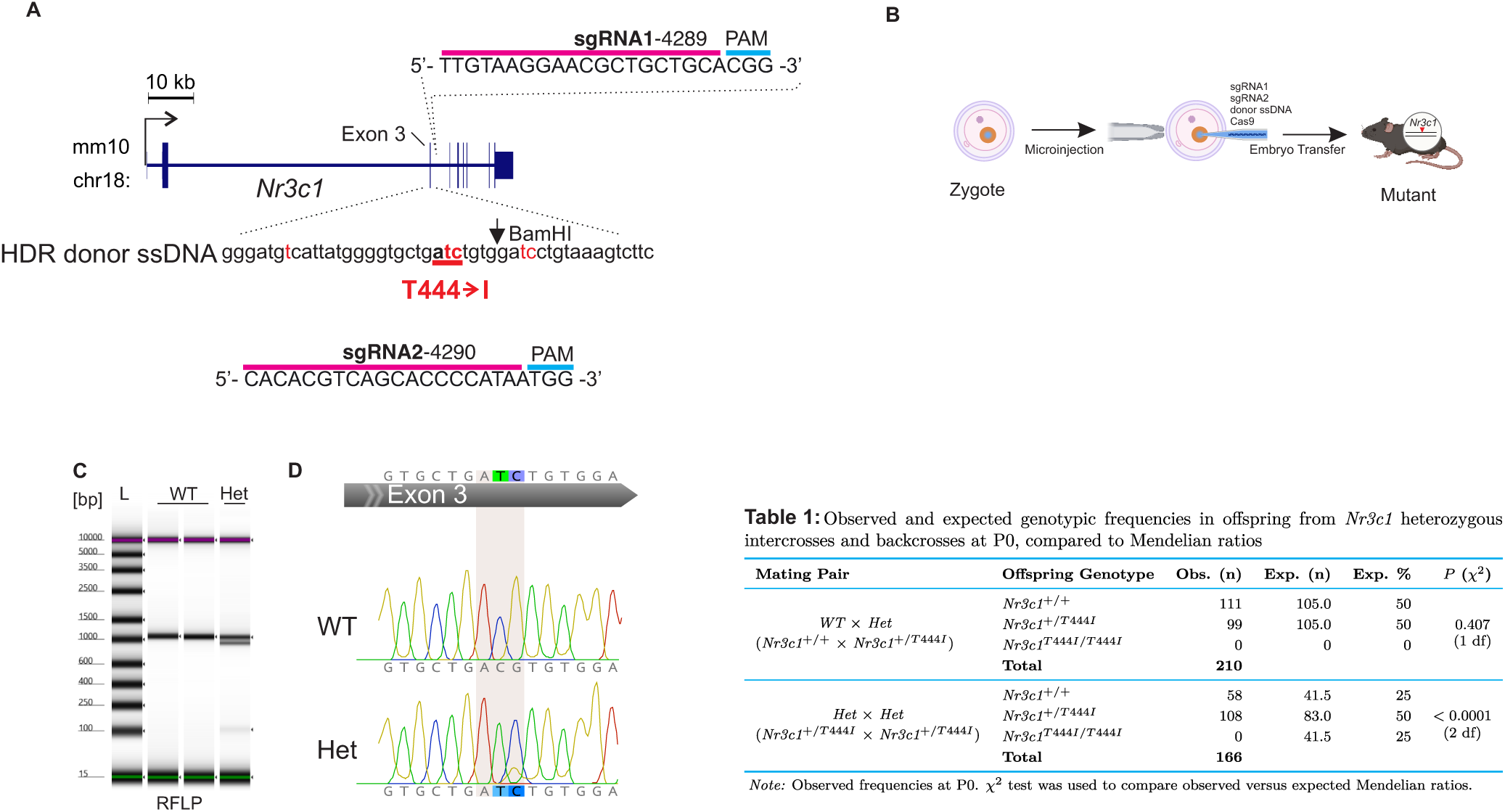
Generation of the Nr3c1-T444I knock-in mouse model using CRISPR-Cas9. (A) Schematic of the gene-editing strategy targeting the *Nr3c1* locus on mouse chromosome 18. Exon 3 was targeted with two single-guide RNAs (sgRNAs). The homology-directed repair (HDR) donor single-stranded DNA (ssDNA) introduced the T444I (Thr444Ile) point mutation (red) and a silent mutation creating a *BamHI* restriction site for genotyping. Scale bar, 10 kb. (B) CRISPR-Cas9 components (sgRNAs, Cas9 protein, and HDR donor ssDNA) were microinjected into zygotes, followed by embryo transfer into pseudopregnant females to generate mutant offspring. (C) Restriction fragment length polymorphism (RFLP) analysis of PCR-amplified products from the targeted region, digested with *BamHI*. Wild-type (WT) samples remain undigested, while heterozygous (Het) samples show digestion products, confirming the T444I mutation. L, DNA ladder. (D) Representative Sanger sequencing chromatograms of exon 3 from WT and Het mice. The WT sequence is unmodified, whereas the Het sequence shows overlapping peaks (shaded region) at the mutation site, indicating both wild-type and mutant alleles. (Table 1) Genotype distributions in offspring from both intercrosses and backcrosses. WT × Het backcrosses (n = 210) yield genotypes consistent with the expected 1:1 ratio of WT and *Nr3c1^+/T444I^* offspring. Het × Het intercrosses (n = 166) deviate from the expected Mendelian 1:2:1 ratio (*P* < 0.0001), with no *Nr3c1^T444I/T444I^* homozygotes observed.

### GR Haploinsufficiency Does Not Result in Overt Tissue Abnormalities

Using the patient-derived GR mutant mouse line (GR-T444I), we performed histological and immunohistochemical analyses to evaluate the in vivo physiological impact of the mutation examined adult heterozygous GR^+/T444I^ (Het) mice (3-6 months old) as well as E12.5 embryos of wild-type (WT), Het, and homozygous GR^T444I/T444I^ (Hom) genotypes. In adult mice, hematoxylin and eosin (H&E) staining of endocrine tissues (pituitary, ovary, and adrenal gland) revealed no detectable morphological differences between Het mutants and WT controls (Fig. 4A). The pituitary glands of both genotypes exhibited a uniform cellular architecture, the ovaries contained healthy follicles, and the adrenal glands showed the expected layered cortical organization. Consistent with the absence of overt histopathology, immunohistochemical staining for cleaved caspase-3 in adrenal gland sections detected no significant difference in apoptosis between Het and WT animals (mean caspase-3 positive cells per section ± s.d.: WT, 0.33 ± 0.17; Het, 0.25 ± 0.17; n = 5 per genotype; P = 0.477; Fig. 4B). Moreover, comprehensive necropsy (gross organ examination and serum chemistry, Extended Data Fig. 4A-N) and histological analysis of other major organs (liver, heart, testes) revealed no signs of toxicity or structural abnormalities in Het mice. Given the critical role of GR in embryogenesis (26, 30, 38) and our observation that homozygous GR^T444I/T444I^ mutants survive to E12.5 but do not reach birth, we next examined mid-gestation embryos. H&E-stained sagittal sections of E12.5 embryos from WT, Het, and Hom litters showed no apparent developmental abnormalities in any major organ system, including the heart, liver, brain, gastrointestinal tract, and limbs (Fig. 4C). All genotypes exhibited GR^T444I/T444I^mutants, embryonic development up to mid-gestation appears grossly normal in these mutants. These findings demonstrate that a single functional GR allele is sufficient to maintain normal organ morphology in adult mice and to support embryonic development up to mid-gestation. The prenatal lethality in Homozygous GR^T444I/T444I^ mutants likely results from impairments in late gestational processes reliant on GR function.

**Figure 4.**
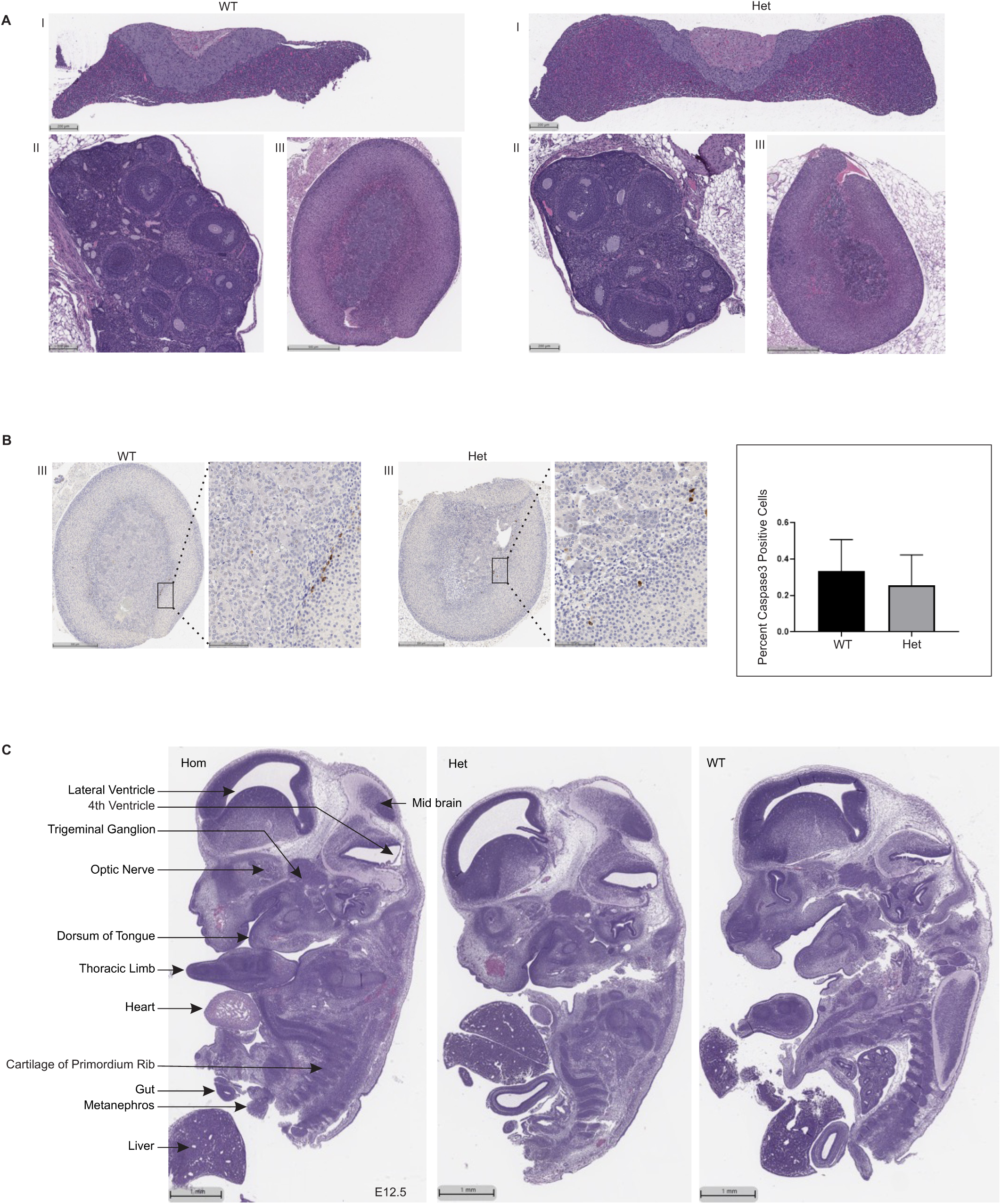
Histological and immunohistochemical analysis of Nr3c1-T444I mutants. (A) Hematoxylin and eosin (H&E) staining of tissue sections from wild-type (WT) and heterozygous (Het) Nr3c1-T444I mice aged 3–6 months. Representative images depict histological evaluation of the pituitary (I), ovary (II), and adrenal gland (III). No observable morphological differences were detected between WT and Het mice. (B) Immunohistochemical (IHC) staining of adrenal glands using an anti-Caspase 3 antibody. Representative images of adrenal sections (III) from WT and Het mice show rare caspase-3-positive cells in both genotypes. Quantification of caspase-3-positive cells was performed using the HALO cytonuclear algorithm. Bar graph represents mean ± s.d. (n = 5 per group), showing no significant difference between WT and Het mice. (C) Sagittal sections of whole embryos at embryonic day 12.5 (E12.5) stained with H&E, comparing wild-type (WT), heterozygous (Het), and homozygous (Hom) *Nr3c1*-T444I embryos. No overt developmental abnormalities were observed in the heart, liver, brain, gut, or limbs in Het and Hom embryos compared to WT. Scale bars are indicated in the images.

### Transcriptomic Analysis Reveals Dosage-Dependent Effects of GR-T444I Mutation on Glucocorticoid Signaling

Given that the GR-T444I mutation results in embryonic lethality during mid-to-late gestation, with no overt structural abnormalities observed at E12.5 under standard histological examination, we hypothesized that this mutation disrupts GR-mediated gene regulation essential for embryonic survival. Supporting this idea, glucocorticoid-stimulated transcription was markedly impaired in proband-derived fibroblasts carrying the orthologous T437I variant. To interrogate this mechanism in vivo, we performed bulk RNA-seq on MEFs from E12.5 GR^WT^, GR^+/T444I^, and GR^T444I/T444I^ embryos treated with 100 nM Dex or vehicle (Fig. 5A). Hierarchical clustering of the 170 differentially expressed, Dex-responsive genes (FDR < 0.05, |log_2_FC| > |log_2_(1.3)|) identified genotype-dependent transcriptional patterns (Fig. 5B,C; Extended Data Fig. 5A,B), with WT cells exhibiting pronounced Dex-induced alterations, heterozygous cells showing partial responses, and homozygous cells displaying minimal reactivity. Median Dex-induced fold changes for up-regulated genes in WT MEFs were attenuated by ∼62% in heterozygotes and ∼92% in homozygotes, with analogous reductions (∼54% and ∼92%, respectively) for down-regulated genes (Fig. 5C; Extended Data Fig. 5A,B). This dosage-sensitive impairment was evident in canonical GR targets critical for signaling and homeostasis, including *Fkbp5*, *Sgk1*, and *Tsc22d3*. Pro-inflammatory genes that are normally repressed by dexamethasone (Dex) in wild-type (WT) cells, including *Ptgs2* and *Ccl7*, exhibited impaired downregulation in cells harboring the T444I mutation. The extent of repression followed a gene dosage-dependent gradient, with heterozygous cells showing an intermediate attenuation and homozygous cells displaying a nearly complete loss of repression. UpSet plots confirmed limited overlap in Dex-responsive genes among genotypes (14 up-regulated genes shared between WT and heterozygotes; Extended Data Fig. 5B), indicating that the mutation not only reduces response amplitude but also reshapes the GR transcriptome. Gene set enrichment analysis of Hallmark pathways corroborated these findings (Extended Data Fig. 5C). Key inflammatory and immune modules—including inflammatory response, TNFα signaling via NF-κB, and Interferon-γ response—were significantly negatively enriched upon Dex treatment in WT and heterozygous GR^+/T444I^ MEFs, but these repressive effects were largely absent in homozygous GR^T444I/T444I^ mutants (Extended Data Fig. 5C). The preserved repression in heterozygous cells, comparable to WT, suggests sufficient GR activity to down-regulate immune programs, which may underlie their developmental viability. Heterozygotes, however, displayed unique negative enrichments of modules including Complement, IL-6–JAK–STAT3 signaling, TGF-β signaling, and apoptosis, hinting at adaptive reprogramming under partial GR loss. Together, these findings support a model in which the T444I mutation impairs GR function in a gene dosage-sensitive manner, leading to progressive attenuation of GR-dependent transcriptional responses to Dex. While heterozygous cells retain partial GR signaling capacity—allowing them to engage both canonical and auxiliary stress-regulatory pathways—homozygous mutants exhibit near-complete transcriptional collapse following Dex exposure, consistent with a functionally null GR allele. These results provide a mechanistic basis for the early embryonic lethality observed in GR^T444I/T444I^ mice.

**Figure 5.**
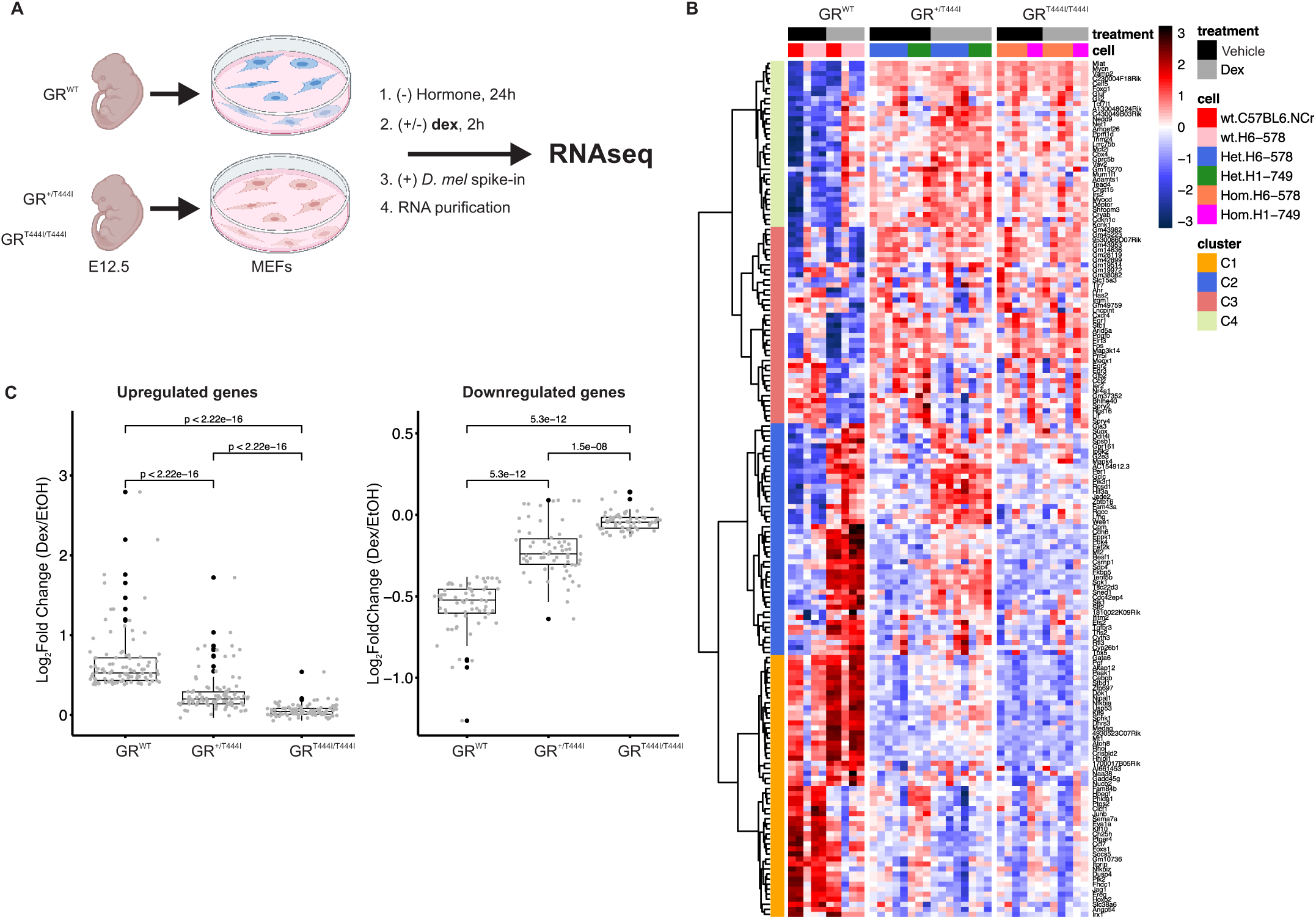
RNA-seq analysis of GR T444I mutant mouse embryonic fibroblasts (MEFs) at E12.5 following dexamethasone (Dex) stimulation. (A) Schematic of the RNA-seq experiment: MEFs from wild-type (GR^WT^), heterozygous (GR^+/T444I^), and homozygous (GR^T444I/T444I^) embryos at E12.5 were cultured in hormone-depleted medium, treated with 100 nM Dex or vehicle (EtOH) for 2 hours, and subjected to RNA extraction with *Drosophila melanogaster* S2 spike-in normalization, followed by sequencing. (B) Heatmap of 170 Dex-responsive genes (union across genotypes; FDR < 0.05, |log₂ FC| > |log₂ (1.3)|), with expression shown as z-scores (low: blue, high: red). Hierarchical clustering (Euclidean distance, Ward.D2) identified 4 clusters: C1 (52 genes), C2 (46 genes), C3 (39 genes), and C4 (33 genes), reflecting distinct expression patterns across treatments and genotypes. Sample annotations are indicated on the heatmap. (C) Boxplots of log_2_fold changes (Dex vs. EtOH) for genes upregulated (left) and downregulated (right) by Dex across the three genotypes. Each point represents an individual gene. Fold changes between genotypes were compared using the paired Wilcoxon rank-sum test, with corresponding *p*-values indicated above the brackets.

### GR-T444I Mutation impairs HPA Axis Regulation leading to Partial Glucocorticoid Resistance in the Mouse Model

Following our characterization of the T444I mutation’s transcriptional and developmental consequences, we evaluated its impact on systemic glucocorticoid signaling in vivo. We examined the hypothalamic-pituitary-adrenal (HPA) axis regulation in adult heterozygous GR^+/T444I^ mice to determine whether this allele mimics glucocorticoid resistance in the proband. To assess basal HPA axis, we measured serum corticosterone—the primary glucocorticoid and main stress hormone in mice—in GR^+/T444I^ and wild-type littermates (aged 3–6 months) at circadian nadir (06:00) and peak (18:00) using enzyme-linked immunosorbent assay (ELISA). Both genotypes maintained a typical diurnal variation of corticosterone, with the dark phase corticosterone levels higher than the light phase levels (Fig. 6A). In the baseline light phase, serum corticosterone levels did not differ significantly between GR^+/T444I^ and WT mice in either sex (males: WT 8.166 ± 2.521 ng/ml, N=3; GR^+/T444I^ 14.081 ± 5.598 ng/ml, N=3; females: WT 27.435 ± 6.345 ng/ml, N=2; GR^+/T444I^ 20.845 ± 0.915 ng/ml, N=2; Fig. 6A). However, in the dark phase, Heterozygous GR^+/T444I^ mice exhibited elevated corticosterone levels as compared to WT controls in both sexes (males: WT 91.07 ± 18.066 ng/ml, N=3; GR^+/T444I^ 167.153 ± 34.612 ng/ml, N=3; females: WT 167.533 ± 8.643 ng/ml, N=3; GR^+/T444I^ 208.85 ± 48.65 ng/ml, N=2; Fig. 6A). These findings indicate dysregulated basal HPA axis activity in heterozygote GR^+/T444I^ mice, consistent with impaired negative feedback control. To further characterize the functionality of the HPA axis, we performed a Dex suppression test, a standard neuroendocrine assay for assessing glucocorticoid sensitivity. Six hours following intraperitoneal administration of Dex, serum corticosterone levels were measured in adult male and female mice (3–6 months old) of both WT and GR^+/T444I^ cohorts. Male WT mice showed significantly lower corticosterone levels than GR^+/T444I^ mice (WT: 48.9 ± 12.6 ng/ml, N=9; GR^+/T444I^: 101.1 ± 13.8 ng/ml, N=9; p=0.013). Female GR^+/T444I^ mice also showed elevated corticosterone levels relative to WT females, but this difference was not statistically significant (WT: 85.7 ± 15.2 ng/ml, N=7; GR^+/T444I^: 123.3 ± 28.0 ng/ml, N=8; p=0.265; Fig. 6B). These data indicate impaired suppression of corticosterone production by dexamethasone in GR^+/T444I^ mice, consistent with partial glucocorticoid resistance, and potential sex-dependent differences. The impaired HPA suppression recapitulates key features of the proband’s clinical phenotype, supporting the role of the T444I mutation in glucocorticoid resistance and associated clinical manifestations.

**Figure 6.**
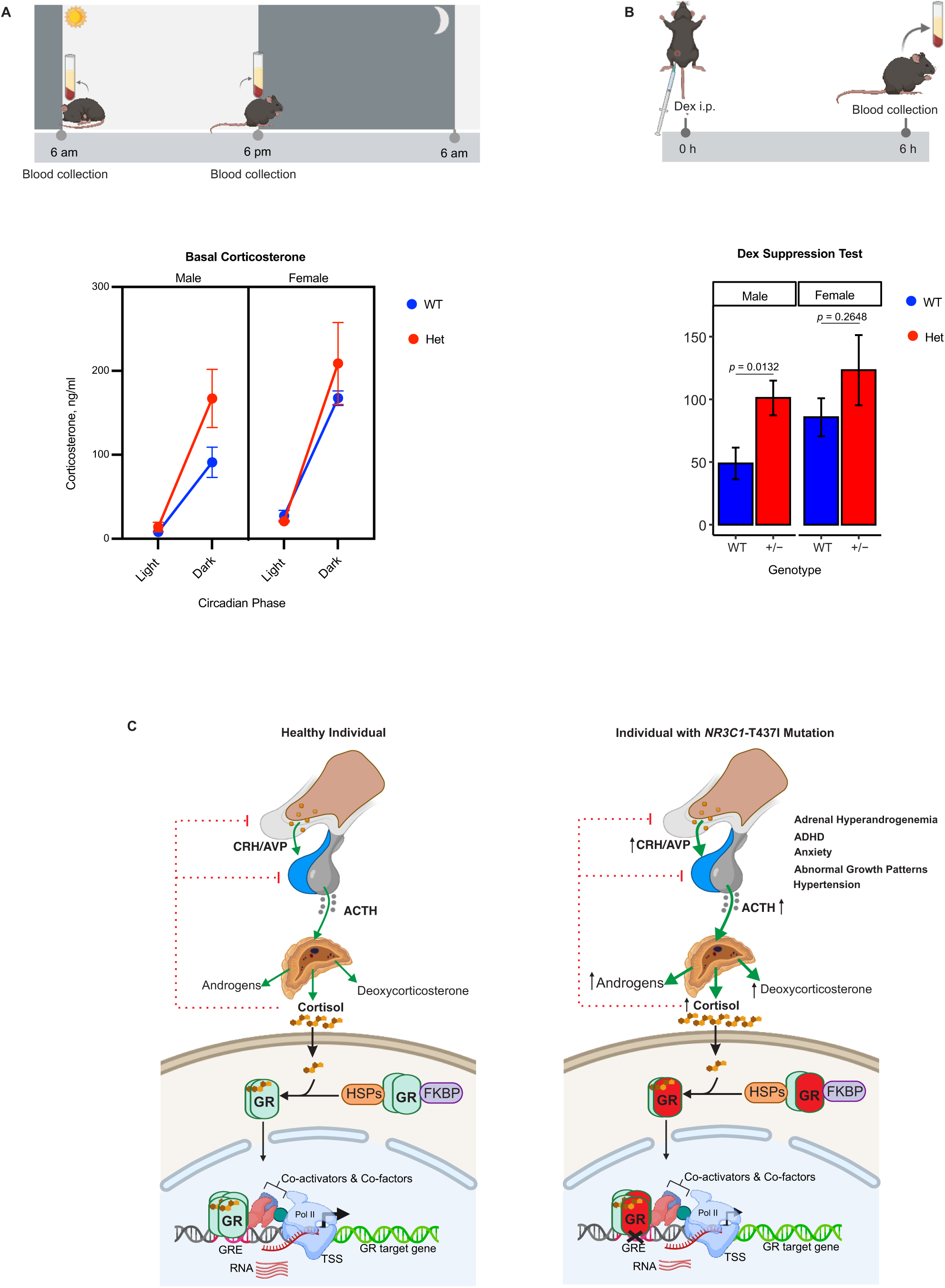
Evaluation of basal corticosterone levels and dexamethasone suppression in WT and GR^+/T444I^ het mice. (A) Schematic of blood collection at 6 am (light phase) and 6 pm (dark phase) for evaluation of basal corticosterone levels. Line graphs display corticosterone concentrations (mean ± s.e.m.) in male and female WT (blue) and Het (red) mice. (B) Schematic of the Dex suppression test. Mice received an intraperitoneal (i.p.) Dex injection at time 0 h, and blood was collected 6 h later. Bar graphs show corticosterone concentrations (mean ± s.e.m.) in male and female WT and Het mice following Dex treatment. **C) A Model for the Pathogenic T437I Mutation’s Impact on GR-Mediated Transcription and Glucocorticoid Signaling**. The figure illustrates the glucocorticoid receptor (GR) signaling pathway in a healthy individual (left panel) compared to an individual with the *NR3C1*-T437I mutation (right panel). (Left Panel -Healthy Individual): In response to hormone stimulation, the hypothalamic-pituitary-adrenal (HPA) axis regulates cortisol production through a negative feedback loop. Corticotropin-releasing hormone (CRH) stimulates the release of adrenocorticotropic hormone (ACTH) from the pituitary gland, which in turn promotes adrenal gland secretion of glucocorticoids, including cortisol. Within target cells, glucocorticoids bind to cytoplasmic GR, leading to its translocation into the nucleus. Once inside, GR binds glucocorticoid response elements (GREs) in the promoter regions of target genes, modulating transcription. This regulation is essential for maintaining homeostasis, metabolism, immune response, and feedback inhibition of the HPA axis, ensuring balanced cortisol levels. (Right Panel – Individual with *NR3C1*-T437I Mutation): In individuals carrying the T437I mutation, the GR retains its ability to translocate into the nucleus upon hormone stimulation. However, the mutation, located in the DNA-binding domain (DBD), potentially impairs GR binding to GREs, leading to a reduced transcriptional response of GR target genes. This disrupted transcriptional regulation weakens the negative feedback control of the HPA axis, resulting in elevated ACTH and cortisol levels. Clinically, this manifests as partial glucocorticoid resistance, contributing to symptoms such as adrenal hyperandrogenemia, ADHD, anxiety, and abnormal growth patterns.

## Discussion

We report the identification and functional characterization of a novel missense variant in *NR3C1* (T437I), located within the DBD of GR, in a family with partial glucocorticoid resistance. The clinical presentation included hypertension and androgen excess, with biochemical evidence of cortisol excess and impaired dexamethasone suppression, characteristic of partial glucocorticoid resistance. To investigate the molecular consequences of this variant in vivo, we generated a knock-in mouse model carrying the orthologous T444I substitution, representing the first patient-derived GR DBD mutation animal model. Heterozygous GR^+/T444I^ mice were viable and fertile, but exhibited elevated basal corticosterone levels and impaired dexamethasone suppression. These endocrine abnormalities closely mirrored key clinical features observed in the patient, supporting the pathogenic relevance of the variant. In contrast, homozygous GR^T444I/T444I^ embryos were non-viable between E12.5 and birth. Although gross morphology appeared normal at E12.5, these embryos ultimately failed to survive gestation, indicating that molecular dysfunction likely precedes the emergence of critical anatomical defects that culminate in embryonic lethality. Supporting this, transcriptomic profiling of MEFs derived from E12.5 embryos revealed a gene dosage–dependent impairment in GR-mediated transcription. Dexamethasone-induced gene expression was reduced by approximately 62% in heterozygous MEFs and nearly abolished, with >90% loss, in homozygous MEFs. These findings demonstrate that the T444I mutation profoundly compromises GR transcriptional output and suggest that failure to activate glucocorticoid-responsive gene programs disrupts essential developmental and endocrine processes (Fig. 6C).

Previous studies of GR-deficient mouse models provide a comparative framework to contextualize the severity of the T444I phenotype. Mice lacking GR entirely (GR^−/−^) survive until E18.5 and die shortly after birth due to lung immaturity and adrenal failure (26). Mice harboring the homozygous GR^dim/dim^ allele, which disrupts DNA binding while preserving partial transrepressive functions, remain viable and fertile (25). Based on these data, it was proposed (25) that GR DNA binding, while critical for full transcriptional activity, is not essential for survival. This “dissociated ligand” model continues to attract support among some investigators. We previously demonstrated that the GR^dim/dim^ mutation does not prevent dimerization, and retains significant DNA binding activity (39, 40). The prenatal lethality of GR^T444I/T444I^ embryos further challenges the original dissociated ligand model and suggests that GR–DNA interactions are indispensable for specific developmental processes that cannot be compensated by residual GR activity alone. The T444I mutation likely disrupts GR function through mechanisms beyond impaired DNA binding, possibly interfering with chromatin recruitment, cofactor assembly, or repression of developmental checkpoint genes. These additional effects may reflect dominant-negative or neomorphic properties that compromise transcriptional networks critical for mid-gestation survival. Although the embryonic lethality of homozygous mutants limits postnatal investigations, and species-specific differences in GR signaling must be considered, this model advances our understanding of GR’s essential roles in early development and challenges long-standing assumptions about the dispensability of DNA binding for receptor function and organismal viability. The concordance between the mouse and human phenotypes highlights the translational relevance of this model, providing a valuable platform for dissecting the mechanisms underlying glucocorticoid resistance and potential therapeutic strategies.

## Methods

### Generation of C57BL/6 *Nr3c1*-T444I Knock-in Mice

#### Design and Synthesis of sgRNAs and ssDNA Donor Carrying the T444I Mutation

The reference sequence for the *Nr3c1* gene was obtained from the University of California, Santa Cruz (UCSC) Genome Browser (transcript: ENSMUST00000115567.7). SgRNAs were designed to target exon 3 using the sgRNA *Scorer 2.0* (41). Five candidate sgRNAs (Supplementary Table 2), IVT-4288, IVT-4289, IVT-4290, IVT-4291, and IVT-4292, were *in vitro* transcribed, complexed with recombinant Cas9 protein, and assessed for genome editing efficiency in mouse embryonic carcinoma P19 cells. Editing efficiencies were determined using high throughput Illumina sequencing (42) (Supplementary Fig. 1). Based on these results, sgRNAs IVT-4289 and IVT-4290 were selected for single stranded DNA (ssDNA) donor design. Chemically modified versions of these sgRNAs were obtained from Synthego for mouse injection experiments. To generate the long ssDNA, first, a double-stranded intermediate plasmid was constructed. DNA containing the T444I mutation and flanking homology arms was synthesized by Twist Biosciences and cloned into the pGMC00018 vector backbone using Gibson assembly (43), yielding the construct pMG0658. This plasmid was then used as a template for long ssDNA production with the Guide-it Long ssDNA Production System v2 (Takara). Primers Nr3c1-ssDNA-F (5ʹ-CTAATTGGCTCTTGCACTGTG-3ʹ) and Nr3c1-ssDNA-R (/5Phos/AATCTAGCCACCACGCTTTG) were used to generate the template for ssDNA synthesis.

#### Pronuclear microinjection of CRISPR/Cas9 complexes into C57BL/6NCr zygotes

All animal procedures were conducted in accordance with NIH guidelines and approved by the Animal Care and Use Committee (ACUC) of the National Cancer Institute (NCI) at Frederick. Female C57BL/6NCr mice (3–4 weeks old) were mated with C57BL/6NCr males to generate fertilized embryos, which were collected from the oviducts at the pronuclear stage for microinjection. To perform genome editing, synthetically modified single-guide RNAs (sgRNAs), IVT-4289 and IVT-4290, were complexed with recombinant Cas9 protein and a single-stranded DNA (ssDNA) donor template. The microinjection solution was prepared with final concentrations of 50 ng/μL for Cas9 protein and 100 ng/μL each for the sgRNAs and ssDNA donor, in a final volume of 30 µL using injection-grade Tris-EDTA (TE) buffer. A total of 1,325 C57BL/6NCr zygotes were microinjected with the CRISPR/Cas9 complexes. After a brief culture period to assess viability, the surviving 622 embryos were surgically transferred into pseudopregnant B6D2F1 females. This procedure resulted in the birth of 74 live pups. Tail biopsies were collected from each pup for genomic DNA extraction, and the presence of the T444I mutation was verified using PCR amplification and sequencing.

#### Genomic screening of Nr3c1-T444I mutant mice using RFLP analysis

To confirm recombination in potential *Nr3c1*-T444I mutant mice, restriction fragment length polymorphism (RFLP) analysis was performed. Genomic DNA was extracted from 74 tail biopsies using a crude extraction method. The tail samples were transferred to a 96-well PCR plate and incubated in 150 µL of a 50 mM NaOH/0.2 mM EDTA solution at 98 °C for 1 h, followed by cooling to 4 °C in a thermocycler. PCR amplification was carried out using Nr3c1-T444I-Screen forward and reverse primers, specifically designed to target the *Nr3c1* gene (see Supplementary Table 3 for primer sequences). The PCR conditions consisted of an initial denaturation at 95 °C for 2 min, followed by 35 cycles of 95 °C for 20 s, 61 °C for 20 s, and 68 °C for 2 min, with a final extension at 68 °C for 5 min. PCR products were digested with *BamHI* (NEB Cat# R0136) and analyzed using the Agilent TapeStation D5000 ScreenTape Assay (Supplementary Information). Successful recombination yielded two fragments of 118 bp and 843 bp, whereas wildtype alleles produced a single 961 bp fragment. For further validation, PCR was performed using Nr3c1-T444I-Seq primers, and the resulting amplicons were purified and subjected to Sanger sequencing to confirm the presence of the T444I mutation. Once mutant founder lines were established, subsequent genotyping was performed using allele-specific primers to distinguish mutant and wild-type alleles (see Supplementary Table 3).

#### Establishing Nr3c1-T444I mouse breeding lines

Founder *Nr3c1*-T444I mice, generated by CRISPR–Cas9 genome editing and confirmed by genotyping, were backcrossed with wild-type C57BL/6NCr mice to obtain F1 offspring. Heterozygous carriers were identified by PCR-based genotyping and used to establish stable breeding colonies. Routine genotyping was performed in subsequent generations to maintain the line. Intercrosses of heterozygous (Het × Het) mice yielded no viable homozygous offspring, prompting timed matings to recover homozygous (Hom) embryos at embryonic day 12.5 (E12.5). For pathological analyses, heterozygous breeding pairs were set up, and successful copulation was confirmed by detection of vaginal plugs. Embryos were harvested by Caesarean section at E12.5 post coitum. Genotyping of all offspring was performed using genomic DNA isolated from tail biopsies.

#### Mendelian segregation analysis of Nr3c1 T444I offspring

Genotypic distribution of offspring was determined by counting the number of animals of each genotype (*Nr3c1^+/+^*, *Nr3c1^+/T444I^*, and *Nr3c1^T444I/T444I^*) from heterozygous intercrosses and control backcrosses. A χ^2^ goodness-of-fit test was used to compare observed frequencies with expected Mendelian ratios. For the control cross (*Nr3c1^+/+^* × *Nr3c1^+/T444I^*; n = 210), counts were compared with the expected 1:1 ratio (χ^2^ = 0.686, df = 1, *P* = 0.407). For the heterozygous intercross (*Nr3c1^+/T444I^* × *Nr3c1^+/T444I^*; n = 166), counts were compared with the expected 1:2:1 ratio (χ^2^ = 55.765, df = 2, *P* < 0.0001). *P* values less than 0.05 were considered statistically significant.

### Neuroendocrinological experiments and analysis

#### Animal Models and Housing Conditions

All procedures were approved by the NIH animal care and use committee and conducted in accordance with national ethical guidelines. Experimental cohorts included wild-type C57BL/6NCr mice and heterozygous (C57BL/6NCr; GR^+/T444I^), males and females. Mice aged 3–6 months were individually housed under a 12:12 h light–dark cycle (lights on at 06:00, lights off at 18:00), with ad libitum access to food and water. All experiments were conducted in the Animal Research Technical Support and Gnotobiotics Core Facility at the Frederick National Laboratory for Cancer Research. Mice were acclimated overnight in the procedure room prior to sampling.

#### Assessment of Basal Corticosterone Rhythms

Circadian corticosterone secretion was assessed using a modified version of the protocol described by Ridder et al(28). Mice were sacrificed at Zeitgeber Time (ZT) 0 (06:00) and ZT12 (18:00), corresponding to the circadian nadir and peak of corticosterone levels, respectively. Animals (n = 6 per genotype per time point) were decapitated without prior anesthesia, and 1 ml of trunk blood was collected. Samples were incubated at room temperature for 15 -30 min, centrifuged at 2,000 × g for 10 min at 4°C, and serum was stored at -80°C. Corticosterone concentrations were quantified using a competitive ELISA (Arbor Assays, Cat# K014-H1) per the manufacturer’s instructions.

#### Dexamethasone Suppression Test

Negative feedback sensitivity of the HPA axis was assessed via intraperitoneal injection of dexamethasone (3 μg/100 g body weight; VEDCO, Cat# VINV-DEX2-100M) administered during the light phase. Mice were euthanized 6 h post-injection for trunk blood collection and corticosterone measurement (n = 12 per genotype).

#### Euthanasia Procedure

Accurate corticosterone assessment requires minimizing stress-induced fluctuations associated with handling. Therefore, euthanasia was performed rapidly by cervical dislocation in awake animals, immediately followed by decapitation using sharp scissors for prompt trunk blood collection.

### Histology and immunohistochemistry

#### Animal care and ethics

All mouse experiments were conducted at the Frederick National Laboratory for Cancer Research, accredited by AAALAC International and compliant with the Public Health Service Policy for the Care and Use of Laboratory Animals. Mouse care adhered to procedures outlined in the “Guide for Care and Use of Laboratory Animals” (National Research Council; 1996; National Academy Press; Washington, D.C.) Guide for Laboratory Animals.

#### Euthanasia and tissue collection

Mice were euthanized by CO2 asphyxiation, following the “Guidelines for the Euthanasia of Mouse and Rats” established by the Animal Care and Use Committee (ACUC) of NCI-Frederick, to minimize pain and suffering. Full necropsy was performed to evaluate the presence or absence of lesions. Terminal blood was collected via cardiac puncture for serum chemistry profiles. Tissues, including the pituitary gland, adrenal glands, and ovaries, were harvested and fixed in 10% neutral buffered formalin (NBF) for histopathological evaluation and immunohistochemistry (IHC) staining.

For embryo collection, pregnant females were euthanized via CO₂ inhalation. A midline laparotomy was performed to expose and remove the uterus, which was immediately transferred to ice-cold phosphate-buffered saline (PBS). Embryos were then harvested by opening the uterine wall and placed in fresh cold PBS for gross inspection. Following examination, embryos were fixed in 10% NBF for downstream histological evaluation.

#### Tissue processing and sectioning

Fixed tissues were processed using an automatic tissue processor and embedded in paraffin blocks. Blocks were sectioned at 5 μm using a manual microtome, and sections were placed on charged slides. Slides were dried at 80°C for 1 hour prior to staining.

#### Hematoxylin and eosin (H&E) staining

H&E staining was conducted using the Sakura Tissue-Tek Prisma automated stainer (Sakura Finetek, Torrance, CA). Slides were hydrated and stained with commercial hematoxylin, clarifier, bluing reagent, and eosin-Y, employing a regressive staining method that overstains tissues followed by a differentiation step to remove excess stain. Slides were cover-slipped using the Sakura Tissue-Tek Glass automatic cover slipper (Sakura Finetek, Torrance, CA) and dried before examination. The histopathological analysis focused on assessing the morphology, consistency, shape, and color of tissues to identify any microscopic lesions or abnormalities

#### Immunohistochemistry (IHC)

IHC for cleaved Caspase-3 was performed using a Leica Bond RX autostainer (Leica Biosystems, Buffalo Grove, IL). Antigen retrieval was achieved with citrate buffer at 100°C for 20 minutes. The primary antibody against Caspase-3 (Cell Signaling Technology #9661, Danvers, MA) was diluted 1:800 and incubated for 60 minutes. Detection was performed using the Bond Polymer Refine Detection kit (Leica Biosystems #DS9800), with the Post-Primary reagent omitted from the default protocol. Normal mouse thymus served as a positive control.

#### Image analysis

Stained slides were scanned at 20× magnification using an Aperio AT2 scanner (Leica Biosystems, Buffalo Grove, IL). Quantitative image analysis was performed using HALO software (v3.6.4134.362; Indica Labs) with the Cytonuclear algorithm (v2.0.9) to determine the proportion of cleaved caspase-3–positive cells. Artifactual regions, including tissue folds and tears, were excluded from analysis. Image annotation and interpretation were performed by a board-certified pathologist (Baktiar Karim).

#### Serum chemistry analysis

At the time of necropsy, terminal blood was collected via cardiac puncture and transferred into heparinized tubes to prevent coagulation. Plasma was isolated by centrifugation and subjected to biochemical analysis using the Vetscan VS2 analyzer (Zoetis Inc., Parsippany, NJ), in accordance with the manufacturer’s protocol. The serum chemistry panel included assays relevant to hepatic and renal function, glucose metabolism, and electrolyte balance. Data were processed and visualized using GraphPad Prism (v10.0; GraphPad Software, San Diego, CA). Statistical comparisons between wild-type (WT) and heterozygous (HET) mice were performed using unpaired two-tailed Student’s t-tests. A *p*-value of <0.05 was considered statistically significant.

### Cell culture

Mouse embryonic fibroblasts (MEFs) and patient-derived skin fibroblasts were maintained in Dulbecco’s Modified Eagle Medium (DMEM; Invitrogen) supplemented with 10% fetal bovine serum (FBS; Gemini Bio-Products) for MEFs and 15% FBS (Gemini Bio-Products) for patient-derived fibroblasts. Both media formulations additionally contained 1 mM sodium pyruvate, 1× non-essential amino acids, and 1× Penicillin–Streptomycin–Glutamine (Gibco; lot no. 223819), corresponding to final concentrations of 0.179 mM penicillin, 0.172 mM streptomycin, and 2.0 mM L-glutamine. Cells were cultured at 37 °C in a humidified incubator with 5% CO₂. For RNA sequencing experiments, fibroblasts were hormone-deprived for 24 h in DMEM supplemented with 10% (MEFs) or 15% (patient-derived fibroblasts) charcoal/dextran-treated FBS, along with 1 mM sodium pyruvate, 1× non-essential amino acids, and 1× Penicillin– Streptomycin–Glutamine. Cells were subsequently stimulated with 100 nM dexamethasone (Dex) for 2 h before RNA isolation.

### Primary Human Fibroblast Isolation

Primary dermal fibroblasts were established from a skin biopsy obtained from the proband II.3. Written informed consent was obtained from the proband’s parents in accordance with the National Institutes of Health Institutional Review Board. For comparison, three primary human dermal fibroblast lines from unaffected donors were obtained from a commercial source (GM01660, AG03512, and GM17592; Coriell Institute for Medical Research, Camden, NJ). Fibroblasts were maintained under the culture conditions described above.

### Isolation of Mouse Embryonic Fibroblasts

Mouse embryos of GR^wt^ (wild-type), GR^+/T444I^ (heterozygous), and GR^T444I/T444I^ (homozygous) genotypes were collected at embryonic day 12.5 (E12.5) and individually minced in 1.5 mL of 0.25% trypsin-EDTA (Gibco) using a sterile blade for 3 minutes. Following incubation at 37°C for 5 minutes, trypsinization was quenched with 4 mL of culture medium. The cell suspension was transferred to a 15-mL tube and allowed to settle for 5 minutes. The supernatant was collected, gently pipetted to obtain a single-cell suspension, and plated onto a 100-mm dish per embryo. Cells were incubated at 37°C in a 5% CO₂ humidified incubator.

### RNA Sequencing

Total RNA was isolated from primary human dermal fibroblasts and mouse embryonic fibroblasts (MEFs) using the PureLink RNA Mini kit (Invitrogen, Thermo Fisher Scientific, 12183018A). A minimum of three biological replicates was included for each condition. To enable normalization, 5% Drosophila S2 cells were spiked into each sample prior to RNA extraction. RNA integrity was assessed with an Agilent Bioanalyzer. For human fibroblasts, strand-specific libraries were prepared from rRNA-depleted total RNA using the TruSeq Stranded Total RNA Library Prep Gold kit (Illumina, 20020598) according to the manufacturer’s instructions. Libraries were sequenced on an Illumina platform by a commercial sequencing provider. For MEFs, strand-specific libraries were prepared with the Illumina Stranded Total RNA Prep, Ligation with Ribo-Zero Plus kit and sequenced at the NIH Intramural Sequencing Center on an Illumina NovaSeq X Plus 1.5B instrument with paired-end reads.

### RNA-Seq Data Processing and Analysis

Raw sequencing reads were demultiplexed into Fastq format using Illumina Bcl2fastq v2.17, permitting up to one mismatch. Adapter sequences and low-quality bases were trimmed with Cutadapt v1.18. The trimmed reads were aligned to the human (hg38) or mouse (mm10) reference genome using STAR(44). Differential expression analysis was conducted using DESeq2(45) (version 1.42.1), which normalized the data by read depth and identified differentially expressed genes (DEGs) between dexamethasone (DEX)-treated and vehicle-treated samples. In parallel, we evaluated normalization using the *Drosophila* spike-in control. However, comparative analyses showed that inclusion or exclusion of the spike-in did not affect global gene expression patterns; therefore, all downstream analyses were conducted using read-depth normalization alone. Log_2_ fold changes and adjusted *p*-values were calculated for each gene, with significance thresholds set at Padj < 0.05 and |log_2_ fold change| > |log_2_(1.3) |.

### Gene set enrichment analysis

GSEA was performed in R (version 4.2.1) using the fgsea package. Ranked gene lists were generated from DESeq2 differential expression analyses, ordering genes by signed log₂ fold-change of Dex-versus vehicle-treated samples. Enrichment was assessed against the MSigDB Hallmark gene set collection (50 curated gene sets). Normalized enrichment scores (NES) and Benjamini–Hochberg–adjusted *P* values were calculated, with significance defined as FDR-adjusted *P* < 0.05. Only pathways reaching this threshold in at least one genotype were included in clustered heatmaps, with asterisks in the figures indicating statistical significance.

## Acknowledgements

This work was supported by the Intramural Research Program of the National Institutes of Health, National Cancer Institute, Center for Cancer Research. We thank Simone Difilippantonio and colleagues for assistance with mouse breeding and dexamethasone suppression testing; Baktiar Karim and team for pathology and immunohistochemistry (supported in part by NIH grant ZIASC010357 to Z.M.Z. and Contract No. HHSN26120150003I); and Roackie Awasthi and colleagues for microinjection of CRISPR reagents. We are also grateful to Raj Chari and his team at the Frederick National Laboratory for Cancer Research for generating CRISPR reagents, and to Dr. Tom Misteli, NIH Distinguished Investigator, for critical review of the manuscript. The content of this publication does not necessarily reflect the views or policies of the U.S. Department of Health and Human Services, nor does mention of trade names, commercial products, or organizations imply endorsement by the U.S. Government.

## Author Contributions

T.T.T. designed and conducted the animal studies, including writing animal study protocols, coordinating the generation of the GR mutant mouse model, performing RFLP analysis, overseeing the dexamethasone suppression test, and carrying out ELISA experiments and analysis, and RNA seq of MEFs. S.V., Q.L., and D.P.M. evaluated and diagnosed the patient and oversaw the genetic analysis. Q.L. and V.K. conducted patient dermal fibroblast derivation and RNA sequencing of proband fibroblast. S.K. and T.T.T. analyzed the RNA sequencing data. L.V. contributed to data interpretation and experimental design. G.L.H. and D.P.M. supervised the project and provided funding. T.T.T. wrote the manuscript with input from S.V., Q.L., S.K., L.V., G.L.H., and D.P.M. All authors reviewed and approved the final manuscript.

---

**Extended Data Fig. 2.**
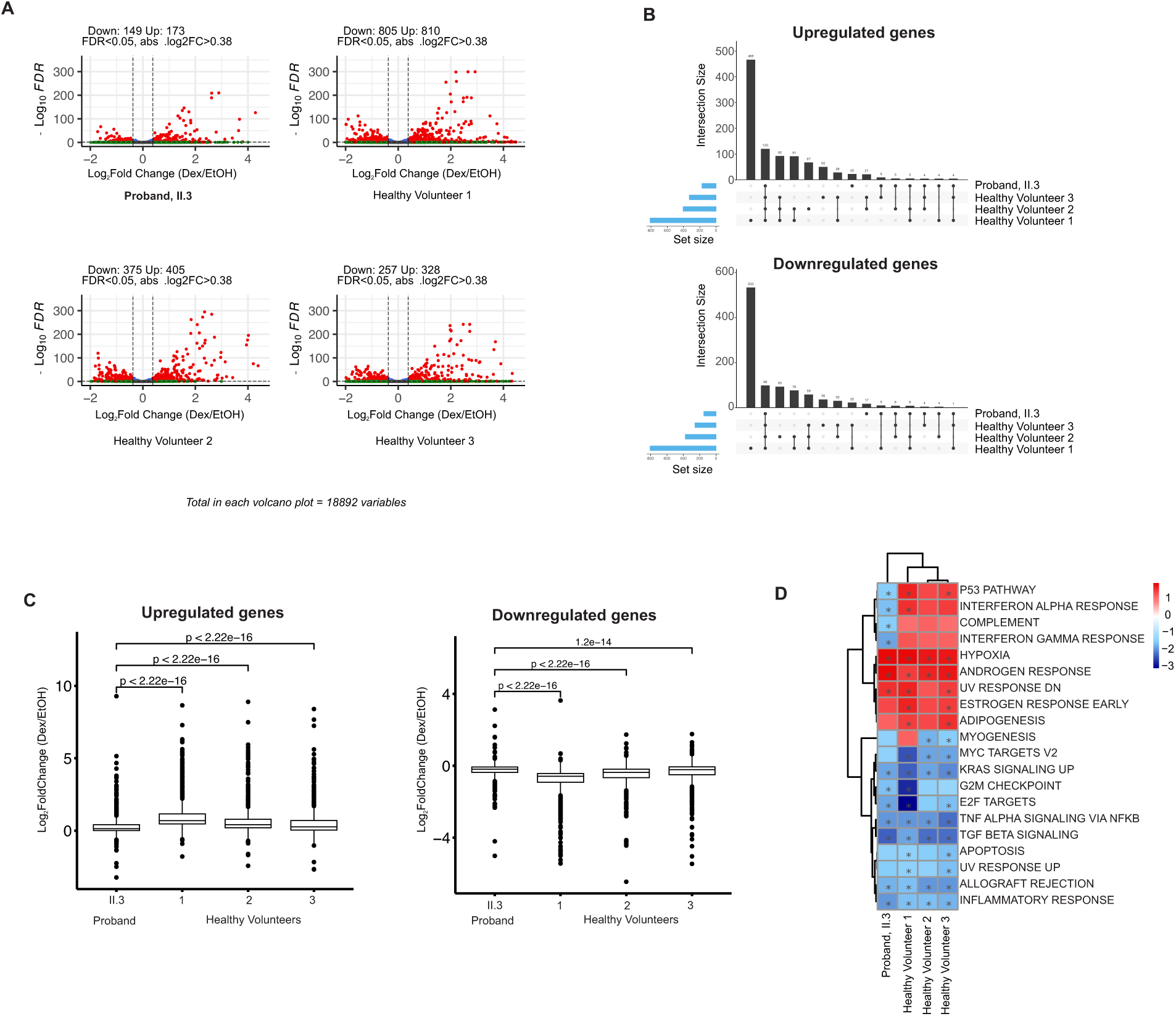
Transcriptomic responses to Dex in proband and healthy volunteers. (A) Volcano plots of differentially expressed genes following Dex treatment in the proband (II.3) and three healthy volunteers. (B) UpSet plots showing unique and shared Dex-responsive genes across individuals. (C) Boxplots showing fold changes of Dex-responsive genes defined in healthy volunteers, plotted across samples from both the proband and the healthy volunteers. (Wilcoxon test; FDR < 0.05; |log_2_FC| > |log_2_(1.3)|) (D) Heatmap of Hallmark gene set enrichment analysis (GSEA) normalized enrichment scores (NES) for pathways significant in the proband or in at least two healthy volunteers (adjusted *P < 0.05*). Asterisks indicate statistically significant pathways (*P < 0.05*).

**Extended Data Fig. 4.**
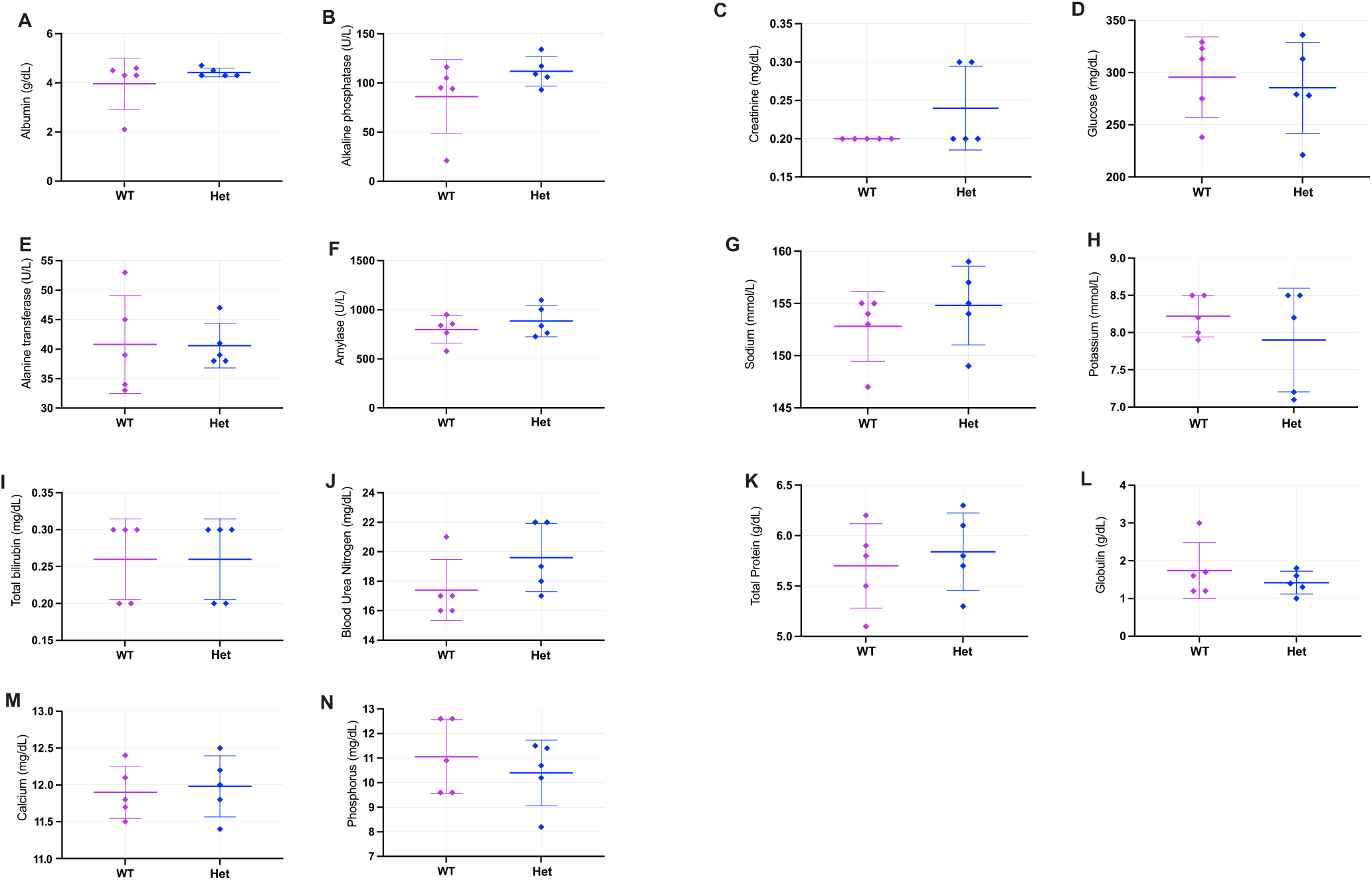
Serum chemistry analyses of *Nr3c1*-T444I mice. Serum biochemical parameters in wild-type (WT) and heterozygous (Het) *Nr3c1*-T444I mice: (A) albumin, (B) alkaline phosphatase, (C) creatinine, (D) glucose, (E) alanine transferase, (F) amylase, (G) sodium, (H) potassium, (I) total bilirubin, (J) blood urea nitrogen, (K) total protein, (L) globulin, (M) calcium and (N) phosphorus. Each point represents one animal; plots show mean ± s.d. No significant differences were detected between WT and Het groups.

**Extended Data Fig. 5.**
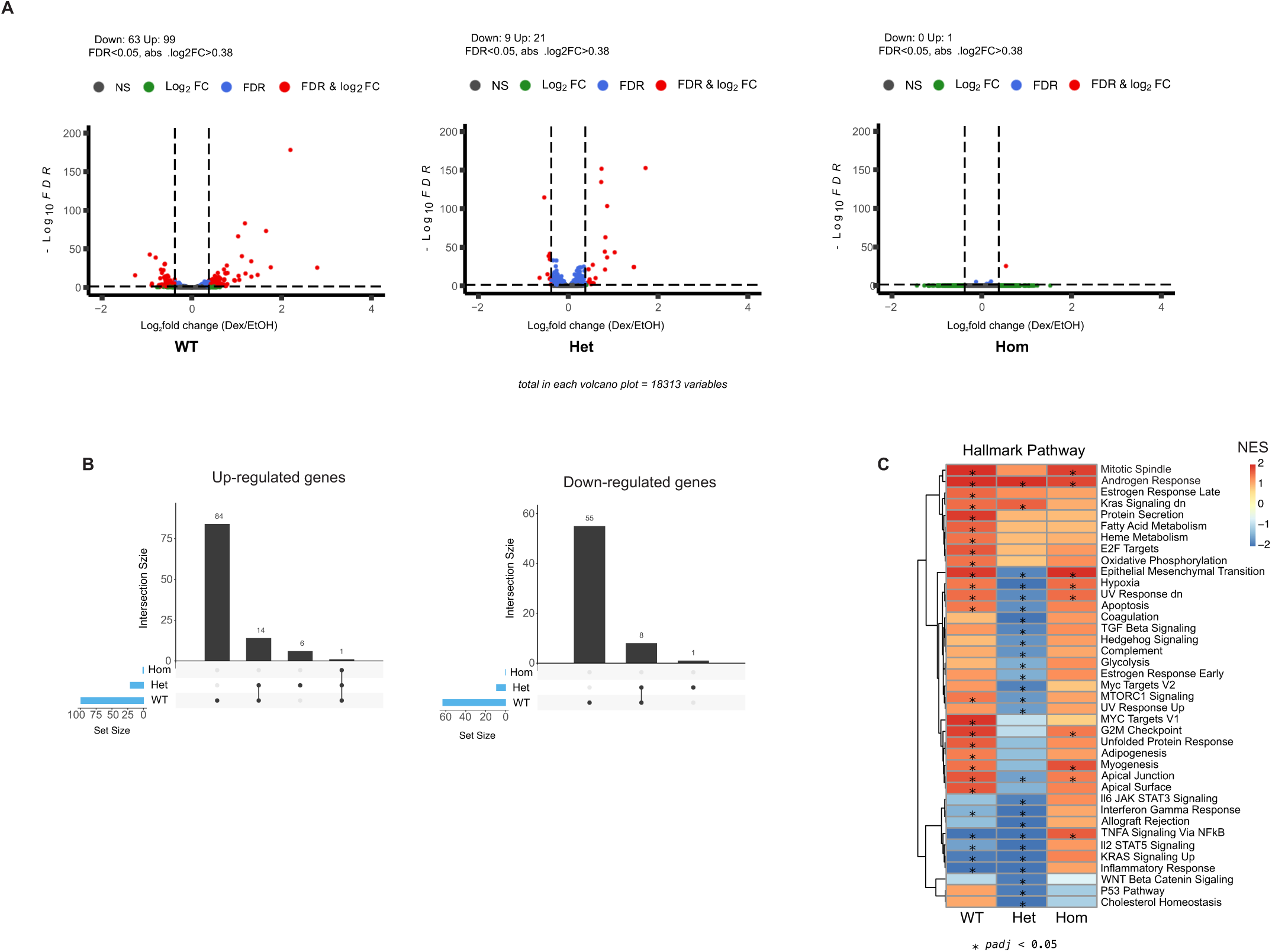
Transcriptomic responses to dexamethasone across *Nr3c1*-T444I MEFs. (A) Volcano plots of differential gene expression following Dex treatment in WT, Het, and Hom MEFs. Differentially expressed genes were defined as FDR < 0.05 and |log₂ fold-change| > |log₂(1.3)|. Analyses were conducted using paired designs: WT (n = 4 pairs), Het (n = 8 pairs), and Hom (n = 6 pairs). (B) UpSet plots showing the overlap and genotype-specific distribution of Dex-responsive genes, displayed separately for upregulated and downregulated sets. (C) Heatmap of normalized enrichment scores (NES) from Hallmark gene set enrichment analysis, including pathways significantly enriched in at least one genotype (adjusted *P* < 0.05). Asterisks denote significant enrichments.

## Supplementary Information

**Supplementary Table 1.**
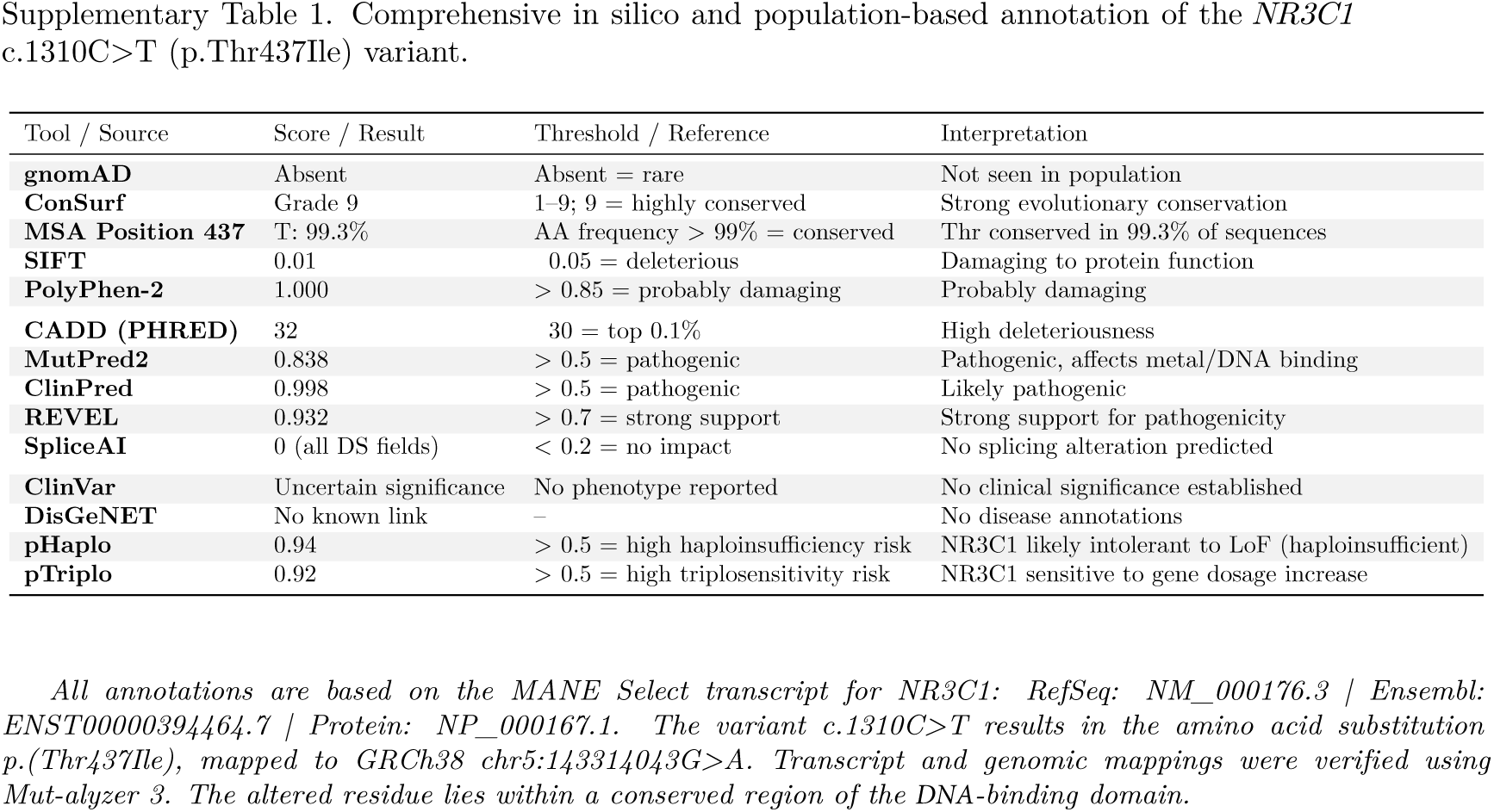
In silico predictions for the *NR3C1* p.Thr437Ile variant. Residue 437 is highly conserved, and multiple computational tools (SIFT, PolyPhen-2, REVEL, CADD, MutPred2, ClinPred) consistently predict a deleterious effect. SpliceAI indicates no splicing impact. The variant is absent from gnomAD, classified as of uncertain significance in ClinVar, and occurs within a conserved region of the DNA-binding domain of *NR3C1*

Comprehensive in silico and population-based annotation of the *NR3C1* c.1310C>T (p.Thr437Ile) variant.
| Tool / Source | Score / Result | Threshold / Reference | Interpretation |
| --- | --- | --- | --- |
| gnomAD | Absent | Absent = rare | Not seen in population |
| ConSurf | Grade 9 | 1-9; 9 = highly conserved | Strong evolutionary conservation |
| MSA Position 437 | T: 99.3% | AA frequency > 99% = conserved | Thr conserved in 99.3% of sequences |
| SIFT | 0.01 | 0.05 = deleterious | Damaging to protein function |
| PolyPhen-2 | 1.000 | > 0.85 = probably damaging | Probably damaging |
| CADD (PHRED) | 32 | 30 = top 0.1% | High deleteriousness |
| MutPred2 | 0.838 | > 0.5 = pathogenic | Pathogenic, affects metal/DNA binding |
| ClinPred | 0.998 | > 0.5 = pathogenic | Likely pathogenic |
| REVEL | 0.932 | > 0.7 = strong support | Strong support for pathogenicity |
| SpliceAI | 0 (all DS fields) | < 0.2 = no impact | No splicing alteration predicted |
| ClinVar | Uncertain significance | No phenotype reported | No clinical significance established |
| DisGeNET | No known link | – | No disease annotations |
| pHaplo | 0.94 | > 0.5 = high haploinsufficiency risk | NR3C1 likely intolerant to LoF (haploinsufficient) |
| pTriplo | 0.92 | > 0.5 = high triplosensitivity risk | NR3C1 sensitive to gene dosage increase |

All annotations are based on the MANE Select transcript for *NR3C1*: RefSeq: NM\_000176.3 | Ensembl: ENST00000394464.7 | Protein: NP\_000167.1. The variant c.1310C>T results in the amino acid substitution p.(Thr437Ile), mapped to GRCh38 chr5:143314043G>A. Transcript and genomic mappings were verified using Mut-alyzer 3. The altered residue lies within a conserved region of the DNA-binding domain.

**Supplementary Table 2.**
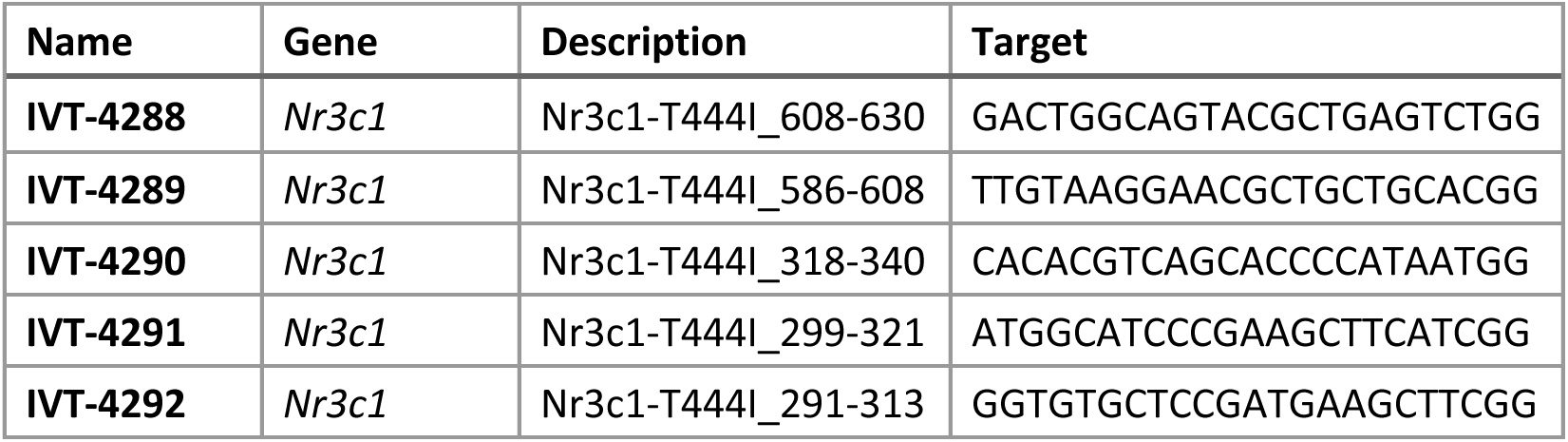
Candidate sgRNAs Designed for Targeting Exon 3 of *Nr3c1* to Introduce the T444I Mutation.

**Supplementary Table 3.**
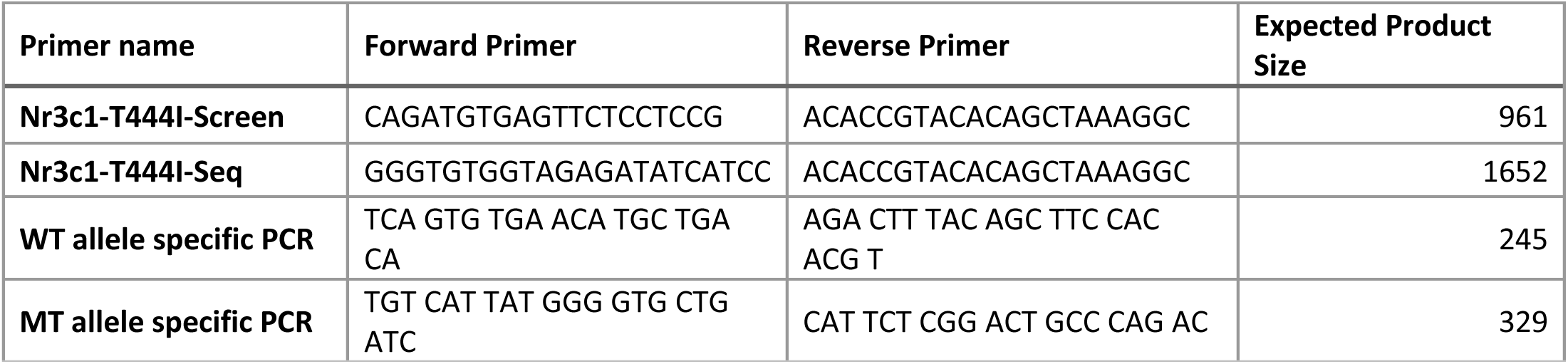
Primer Sequences Used for Genotyping of Nr3c1-T444I Mutant Mice.

**Supplementary Figure 1.**
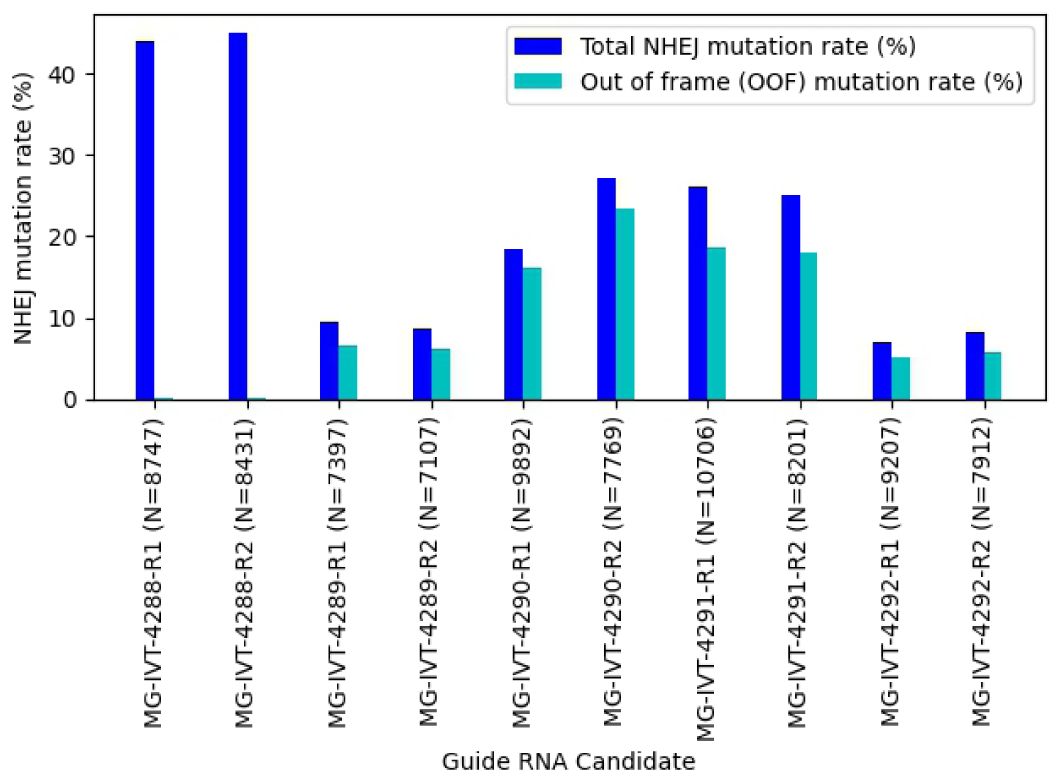
Genome editing efficiency of sgRNA candidates targeting *Nr3c1* exon 3. Non-homologous end joining (NHEJ) mutation rates (blue) and out-of-frame (OOF) mutation rates (cyan) were measured in P19 cells following CRISPR/Cas9 editing with five candidate sgRNAs. The number of sequencing reads (N) for each guide RNA replicate is shown on the x-axis. Based on these results, sgRNAs IVT-4289 and IVT-4290 were selected for subsequent donor design and mouse microinjection experiments.

## Statistical information

Statistical analyses were performed using GraphPad Prism (version 9.0) or R (version 4.2.1), with Bioconductor packages used for advanced data processing and visualization. Appropriate pairwise comparisons used to calculate *P* values are specified in the respective figures and figure legends. Statistical significance was defined as *P* < 0.05. Data are presented as mean ± s.e.m., with the number of biological replicates (*N*) indicated in each figure legend. Outliers were excluded using the 1.5 × interquartile range (IQR) rule.

## Data availability

RNA-seq data generated in this study have been deposited in the NCBI Gene Expression Omnibus (GEO) under accession number GSE305914. All other relevant data are available from the corresponding author upon reasonable request. Source data for the figures are provided in the supplementary information accompanying this paper.

